# Rapid genomic convergent evolution in experimental populations of Trinidadian guppies (*Poecilia reticulata*)

**DOI:** 10.1101/2021.02.10.430609

**Authors:** Mijke J. van der Zee, James R. Whiting, Josephine R. Paris, Ron D. Bassar, Joseph Travis, Detlef Weigel, David N. Reznick, Bonnie A. Fraser

**Affiliations:** Biosciences, University of Exeter, Stocker Road, Exeter, EX4 4QD, UK; Department of Biology, Williams College, Williamstown, MA, USA; Department of Biological Science, Florida State University, Tallahassee, FL, USA; Department of Molecular Biology, Max Planck Institute for Developmental Biology, Tübingen, Germany, 72076; Department of Biology, University of California Riverside, Riverside CA, USA, 9252120

**Keywords:** Experimental evolution, Convergent evolution, Rapid evolution, Population genomics, guppies, Poecilia reticulata

## Abstract

It is now accepted that phenotypic evolution can occur quickly but the genetic basis of rapid adaptation to natural environments is largely unknown in multicellular organisms. Population genomic studies of experimental populations of Trinidadian guppies (*Poecilia reticulata*) provide a unique opportunity to study this phenomenon. Guppy populations that were transplanted from high-predation (HP) to low-predation (LP) environments have been shown to mimic naturally-colonised LP populations phenotypically in as few as 8 generations. The new phenotypes persist in subsequent generations in lab environments, indicating their high heritability. Here, we compared whole genome variation in four populations recently introduced into LP sites along with the corresponding HP source population. We examined genome-wide patterns of genetic variation to estimate past demography, and uncovered signatures of selection with a combination of genome scans and a novel multivariate approach based on allele frequency change vectors. We were able to identify a limited number of candidate loci for convergent evolution across the genome. In particular, we found a region on chromosome 15 under strong selection in three of the four populations, with our multivariate approach revealing subtle parallel changes in allele frequency in all four populations across this region. Investigating patterns of genome-wide selection in this uniquely replicated experiment offers remarkable insight into the mechanisms underlying rapid adaptation, providing a basis for comparison with other species and populations experiencing rapidly changing environments.

**IMPACT STATEMENT:** The genetic basis of rapid adaptation to new environments is largely unknown. Here we take advantage of a unique replicated experiment in the wild, where guppies from a high predation source were introduced into four low predation localities. Previous reports document census size fluctuations and rapid phenotypic evolution in these populations. We used genome-wide sequencing to understand past demography and selection. We detected clear signals of population growth and bottlenecks at the genome-wide level matching known census population data changes. We then identified candidate regions of selection across the genome, some of which were shared between populations. In particular, using a novel multivariate method, we identified parallel allele frequency change at a strong candidate locus for adaptation to low predation. These results and methods will be of use to those studying evolution at a recent, ecological timescale.

## INTRODUCTION

Historically, evolution in natural populations was thought to happen on a long timescale, one that could not be observed in real time (Gillespie 1994). However, recently it has become clear that evolutionary change can be so rapid it occurs on an ecological timescale and many studies have now shown that phenotypic traits can evolve within a few generations (Endler 1980; Grant and Grant 2020; Losos 2011). Understanding the genetic basis of rapid adaptation however, is still in its infancy. Studying the effect of environmental changes at the genomic level provides insight into genetic constraints and mechanisms of evolutionary adaptation (Pascoal *et al.* 2014), which in turn could help develop conservation efforts tailored to a specific species or population, and ultimately help predict a population’s response to future changes in their environment.

Until recently it has been difficult to detect recent genomic adaptation. Firstly, rapid genetic adaptation is likely to occur through soft sweeps on standing genetic variation (SGV) because adaptation is most rapid when the beneficial alleles are already present in the populations, whereas in hard sweeps it takes time for beneficial *de novo* mutations to appear (Barrett and Schluter 2008; Hermisson and Pennings 2005). In a soft sweep, multiple haplotypes can rise to high frequency and as a result, the effect of selection on levels of diversity and the allele frequency distribution are usually less severe, making them difficult to detect (Garud *et al.* 2015). Secondly, whole genome sequencing (WGS) for non-model organisms is limited, and reduced representation approaches (such as microsatellites and RAD-seq) only sample a small proportion of the genome and cannot be expected to detect loci involved in adaptation (Lowry *et al.* 2017; Tiffin and Ross-Ibarra 2014). Now, with the accessibility of WGS for non-model organisms and improved methods for detecting soft sweeps, we can begin to address this long-standing problem in evolutionary biology.

Unfortunately, distinguishing genomic signatures of selection from random genomic changes can be difficult. Populations that have evolved convergent phenotypes in response to similar environmental changes provide an opportunity to leverage signatures of repeated selection to distinguish false positives due to drift from true positives. The occurrence of convergent evolution is considered strong evidence for natural selection, as processes other than selection (such as genetic drift and random mutations) are unlikely to result in the same evolutionary changes in independent populations. Here, following Lee and Coop (2017), we use the term convergence to define the repeated evolution of the same allele in independent populations, which includes both classically defined convergent and parallel evolution. By looking for signatures of convergent evolution in rapidly evolving, replicated populations, we should be able to distinguish random genomic changes from those caused by selection.

### The Trinidadian guppy system

The guppy system in the Northern Range Mountains of Trinidad is a well-known model for studying phenotypic evolution in wild and experimental populations, offering a powerful platform to investigate convergent genomic adaptation. The mountains are drained by a set of parallel rivers punctuated by waterfalls preventing upstream colonisation by larger fish species, including the guppy predators *Crenicichla alta* and *Hoplias malabaricus* (Magurran 2005). Above the waterfalls there is a relatively low-predation (LP) environment, whereas the drainages downstream of the barriers are considered a high-predation (HP) environment. Males from LP environments are more colourful, have a larger number of spots and larger individual spots than their HP counterparts (Endler 1980). Additionally, male and female LP guppies have larger body sizes at maturation and reproduce less frequently than guppies occupying HP environments (Reznick and Bryga 1996; Reznick and Endler 1982). This pattern is broadly replicated in rivers across the Northern Range Mountains, but there is variation in some aspects of phenotypes from similar environments (Endler and Houde 1995; Kemp *et al.* 2009). Many of the phenotypic differences between HP and LP populations are also heritable in laboratory experiments (Reznick and Bryga 1996; Reznick and Endler 1982), suggesting convergent phenotypic evolution has a genetic basis.

A unique opportunity to investigate rapid adaptation is provided by four replicated experimental populations of guppies in the Northern Range Mountains of Trinidad. In 2008 and 2009, guppies were transplanted from a single high predation (HP) source in the Guanapo river (“GHP”) to four, previously guppy-free, low predation (LP) environments: Lower Lalaja (“ILL”), Upper Lalaja (“IUL”), Caigual (“IC”) and Taylor (“IT”) (figure 1a). Additionally, in each introduction year pair, the canopy of one of the localities was thinned (IUL and IT in for 2008 and 2009, respectively, figure 1a), which increased the primary productivity of these streams (Kohler *et al.* 2012). In 2008, 38 fish of each sex were introduced to ILL and 38 to IUL. In March 2009, 52 fish of each sex were introduced into IT and 64 fish of each sex to IC (Arendt *et al.* 2014). The methods of sampling from the wild population and introducing fish to the new localities were the same for each population and have been described by Travis et al. (2014). Briefly, juveniles sampled from GHP were raised to maturity in single-sex groups, then housed in tanks of 5 males and 5 females for approximately three weeks to enable mating before introduction. Females of each tank were randomly assigned to one of the paired streams (e.g. IUL in the first pair), while the males of the tank were assigned to the opposite pair (e.g. ILL in the first pair). The consequence is that all males were represented in both streams, either as sires of embryos or stored sperm from mating that occurred before introduction or by being introduced into that stream, thus increasing the effective population size relative to census size. Importantly, the HP fish introduced each year were collected in that year making the stream pairs initiated on separate years independent from each other.

**Figure 1.**
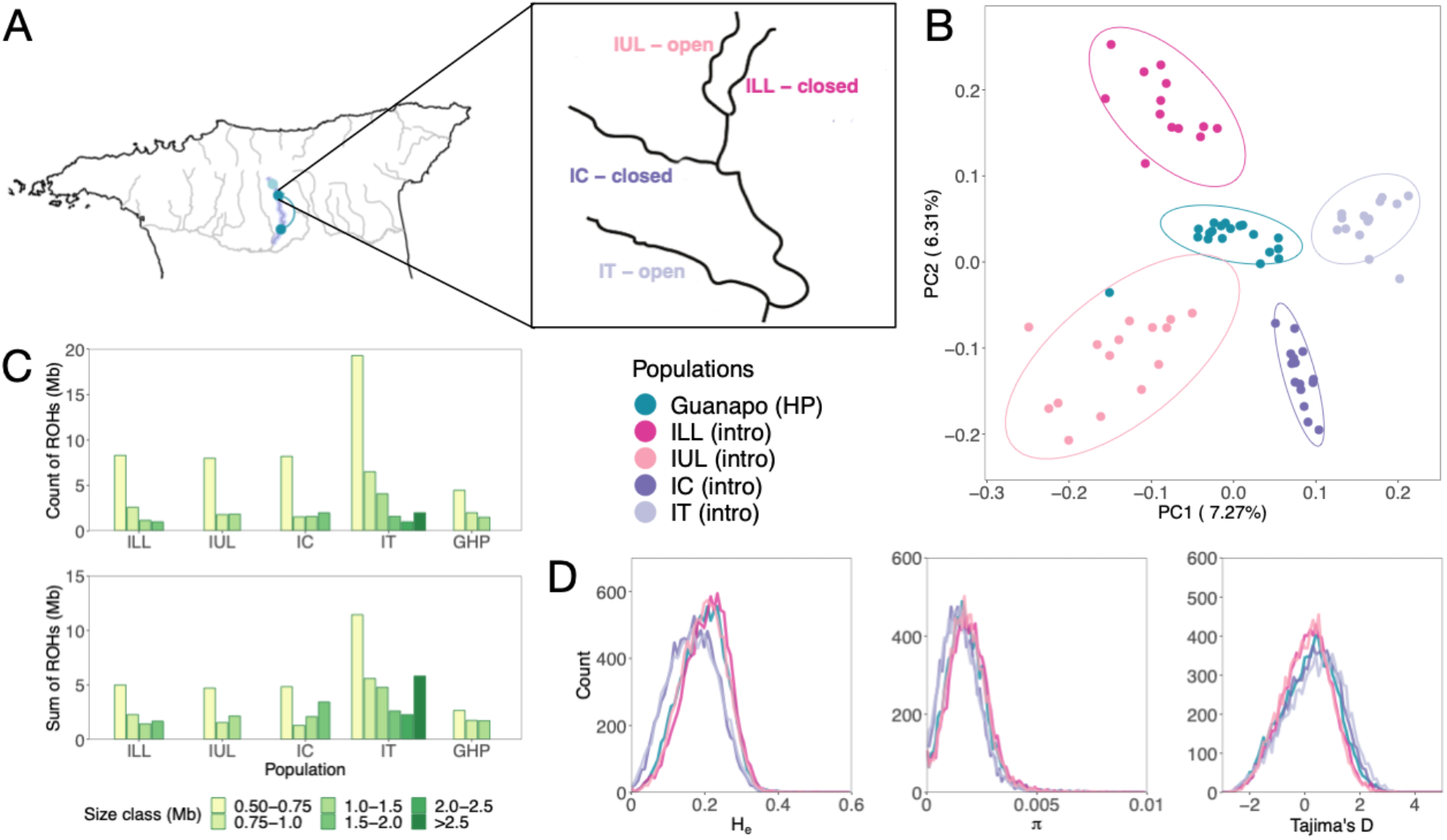
Sampling sites, population structure, runs of homozygosity and neutral statistics across the introduction sites and the HP source. Map (A) highlights the Guanapo river with the sampled populations, and an inset shows the experimental rivers. “Open” indicates sites with manipulated canopies, “closed” indicates intact canopies. PCA (B) with populations coloured according to river. Number and sum of runs of homozygosity (ROH) (C) in different size classes, per population. Expected heterozygosity (He), nucleotide diversity (π) and Tajima’s D (D) for each of the introduced populations and their HP source.

Phenotypic evolution in these populations has been well documented. Kemp et al. (2018) found that all introduced populations, except for IC, increased in coverage of blue/green spots compared to the source population, within 8-12 generations (4-6 years since introduction, assuming a generation time of 2 per year). In IUL and IT, there was also a reduction in black spots. After 4-6 generations, introduced males matured at a later age and larger size in all 4 populations. This evolution became evident only after populations had reached their peak densities, which differed between the introduction years (Reznick *et al.* 2019). ILL and IUL were introduced with much lower initial densities (due to fewer introduced fish and larger stream size than IC and IT) and it took 3 years for them to attain peak density. IC and IT, on the other hand, had high initial densities and reached peak densities after just 2 years (Reznick *et al.* 2019).

In addition to overall population growth, all four populations experienced seasonal fluctuations, with populations decreasing during the rainy seasons. Population densities under thinned canopies (IUL and IT) fluctuated more than under intact canopies (ILL and IC). These fluctuations were particularly strong in IT, where Reznick et al. (2019) observed the near extinction of the population during the rainy season in the first year. Finally, 6-8 generations after the introduction, all populations had acquired a plastic response in growth rate and resting metabolic rate to predator cues in an otherwise common lab environment (Handelsman *et al.* 2013).

Here we capitalise on this unique set of rapidly evolving experimental populations to investigate the early stages of adaptation. We first examine patterns of genetic variation with whole genome sequence data to understand the impacts that neutral processes may have had on the populations. Then, we investigate signals of selection across the genome and instances of genomic convergence amongst the four populations with a combination of selection scans and a multivariate analysis of allele frequency change vectors.

## METHODS

### Sampling, sequencing and SNP calling

Fish from the four experimental populations and ancestral population were sampled in the spring of 2013, which is 2 years (3-5 generations) after the evolution of male age and size at maturity described in Reznick et al. (2019) and 4-5 years after the introduction. We collected approximately 20 individuals from each site (total N=94, both male and females, table S1). Each sample was sequenced across the genome at greater than 7X coverage. Reads were mapped to the updated guppy genome (Fraser *et al.* 2020), SNPs were called using GATK (v4.0.5.1), and were then phased following (Malinsky *et al.* 2018). For more details see supplemental information.

### Genome-wide diversity, population structure and runs of homozygosity

Principal Component Analysis (PCA) was performed on a linkage-pruned VCF (-- indep-pairwise 50 5 0.2) in Plink v1.90b6.7 (Chang *et al.* 2015) to assess population structure. Population specific summary statistics were calculated with PopGenome (nucleotide diversity (***π***), Tajima’s D and global F_ST_) (Pfeifer *et al.* 2014) and VCFtools v0.1.16 (expected heterozygosity, H_e_)(Danecek *et al.* 2011). Runs of homozygosity (ROH) were calculated for each individual with a sliding window approach with 50 SNPs per window using Plink.

### Selection scans

Allele frequencies were calculated in VCFtools to investigate how many fixed differences exist between GHP and each LP population, as well as allele frequency changes (ΔAF) between GHP and each of the LP populations. This was done by calculating the change in minor allele frequency in non-overlapping sliding windows of 75,000 bp. The introduced populations have only recently been established and we found only slight differences in divergence between these populations and the GHP source (see results). Therefore, we chose to use two statistics based on haplotype homozygosity that are more suitable for detecting very recent directional selection: cross-population extended haplotype homozygosity (XP-EHH), which was developed to detect selective sweeps that are nearly fixed in one population but still polymorphic in the population as a whole (Sabeti *et al.* 2007) and iHH12, which combines the top two most frequent haplotypes into a single haplotype, making it more effective at detecting soft sweeps (Garud *et al.* 2015; Torres *et al.* 2018). More details can be found in supplemental methods.

### Multivariate analysis of parallel allele frequency changes

We adapted De Lisle and Bolnick’s (De Lisle and Bolnick 2020) multivariate approach for interpreting parallel evolutionary change to investigate parallel allele frequency changes in all four populations simultaneously. Instead of using phenotypic change vectors, we applied the method to vectors of allele frequency change between the HP source and each of the introduced populations. Briefly, we divided each chromosome into non-overlapping windows of 200 SNPs, and calculated a matrix of normalised allele frequency changes per window:

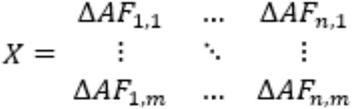

Where *n* is the number of SNPs in a window and *m* the number of populations. For each of these matrices we then computed the correlation matrix describing correlations of normalised allele frequencies among the vectors of each HP-LP pair, by multiplying matrix *X* with its transpose:

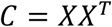

Eigen decomposition of each C matrix produces a distribution of eigenvalues; distributions with excessive variance explained by the first eigenvector suggest a shared direction of allele frequency changes among all population pairs through multivariate space. This direction of change could be parallel (along the same axis in the same direction) or anti-parallel (along the same axis in different directions). We then created a null-distribution to determine which windows show extreme eigenvalues.

## RESULTS

### Population structure and genome-wide diversity reveal limited divergence between the experimental populations and their source

PCA revealed the introduced populations to have only slightly diverged from the GHP source. PC1 and PC2 accounted for 7.3% and 6.3% of the variation, respectively (figure 1b); PC1 separates the introduction years from each other, while PC2 separates the introduction populations in different directions. PC3 and PC4 did not further separate populations and all other PCs accounted for <5% of the variation (figure S1). When compared to natural populations across the Northern Range described in Whiting et al. (2020), the experimental populations show very limited divergence from their source (GHP) (figure S2). Pairwise F_ST_ was low between introduced populations and their source (median F_ST_ = 0.013 – 0.023), with the strongest difference found between IT and GHP (table S2). This higher F_ST_ between IT and GHP, despite the observation of relatively low divergence between the two on PC2, can be explained by the low within-population variation of IT, resulting in GHP variants that are more likely to have been lost in IT. Recall that IT experienced a population crash during the first wet season.

Among experimental populations, IC, IUL, and ILL were more distinct from IT than they were from one another (table S2). Guppies from populations initiated in the same year were more similar to one another than guppies from populations initiated in other years (table S2). This is because there were two independent collections of GHP fish, one per introduced pair, and the introductions were designed to introduce the same distribution of genetic variation into each member of a pair.

The introduction of guppies from GHP to ILL, IUL, IC and IT led to very minor changes in genetic diversity within the introduction populations (figure 1d, table S3). IT experienced a decrease in H_e_ of 15.8% and IC a 15.5% decrease (p<0.0001 for all three, Mann-Whitney U test), whereas ILL and IUL experienced very slight increases of H_e_ relative to GHP: 0.5% in IUL (p = 0.07 Mann-Whitney U test) and 5.4% in ILL (p < 0.0001). Similarly, median nucleotide diversity (π) decreased by 15.1% in IT and 14.4% in IC (p<0.0001, for both), and was slightly, but significantly, increased in ILL and IUL (8.0% and 2.6%, p<0.0001 and p=0.002, respectively, Mann-Whitney U test). Tajima’s D was shifted to a more positive value in IC and IT compared to GHP (figure 1d, table 3, p<0.0001). This suggests a loss of rare alleles compared to the HP source, which is consistent with a sudden population contraction. In ILL and IUL Tajima’s D has decreased slightly compared to GHP, but stayed positive (figure 1d, table S3, p<0.0001), implying an increase of rare alleles which could be related to a recent selective sweep or rapid population expansion.

### Runs of homozygosity provide evidence of recent inbreeding and a bottleneck in IT

We found ROH of > 0.5 Mb in 93 of the 94 individuals, with one individual in GHP lacking the ROH fitting requirements we set *a priori*. Of the four introduced populations, IT had nearly three times more ROH than the other introduced populations (N=423, table S4), which had numbers comparable among themselves (IUL (N=151), IC (N=150) and ILL (N=142)). GHP had the lowest number of ROH (N=83). This pattern is again consistent with IT having experienced a population crash during the first year of the experiment.

We also observed variation in the maximum length of ROH among populations. IT was the only population with ROH > 2Mb (figure 1c, table S4), suggesting that this population experienced recent inbreeding. ILL, IUL, and IC have more ROH and a larger proportion of their genome covered in ROH compared to GHP, suggesting there may have been a limited bottleneck effect since the introduction, although not as severe as that in IT (figure 1c).

Inbreeding coefficients (F_ROH_) were significantly higher in IT compared to the other introduced populations (table S4, p<0.0001, Mann-Whitney U test), again confirming recent inbreeding in this population. The remaining three introduction populations did not significantly differ from each other (ILL-IC: p=0.274, ILL-IUL: p=0.650 and IUL-IC: p=0.624, Mann-Whitney U test), but did have slightly higher values compared to GHP (GHP-ILL: p<0.0001, GHP-IUL: p=0.002 and GHP-IC: p=0.002, Mann-Whitney U test).

### There are strong signals of selection in all four populations despite limited divergence genome-wide

We found little change in allele frequency in the introduced populations. Mean allele frequency change was low in all populations (table S5). There were no fixed differences in allele frequency between GHP and any of the four introduction populations. Of the SNPs that were the minor allele in GHP, fewer became fixed in ILL and IUL than in IC and IT (N=97 in IIL, N=102 in IUL, N=385 in IC, and N=469 IT, table S5). None of these fixed SNPs were shared among all four populations. IC and IT had more sites than ILL and IUL where allele frequencies had not diverged from GHP, consistent with the shorter period of growth they experienced before the onset of selection in these populations (Reznick *et al.* 2019).

We analyzed 9,804 windows of 75kb length using XP-EHH and iHH12 to evaluate regions under selection in the experimental populations. For the XP-EHH statistic, IT had the most outlier windows (N=50), followed by ILL (N=39), IUL (N=17) and finally IC (N=14) (figure 2a, tables S6-S9). For iHH12, IT again had the most outliers (N=41), followed by ILL (N=39), IC (N=36), and IUL (N=19) (figure 2b, tables S10-S13).

**Figure 2.**
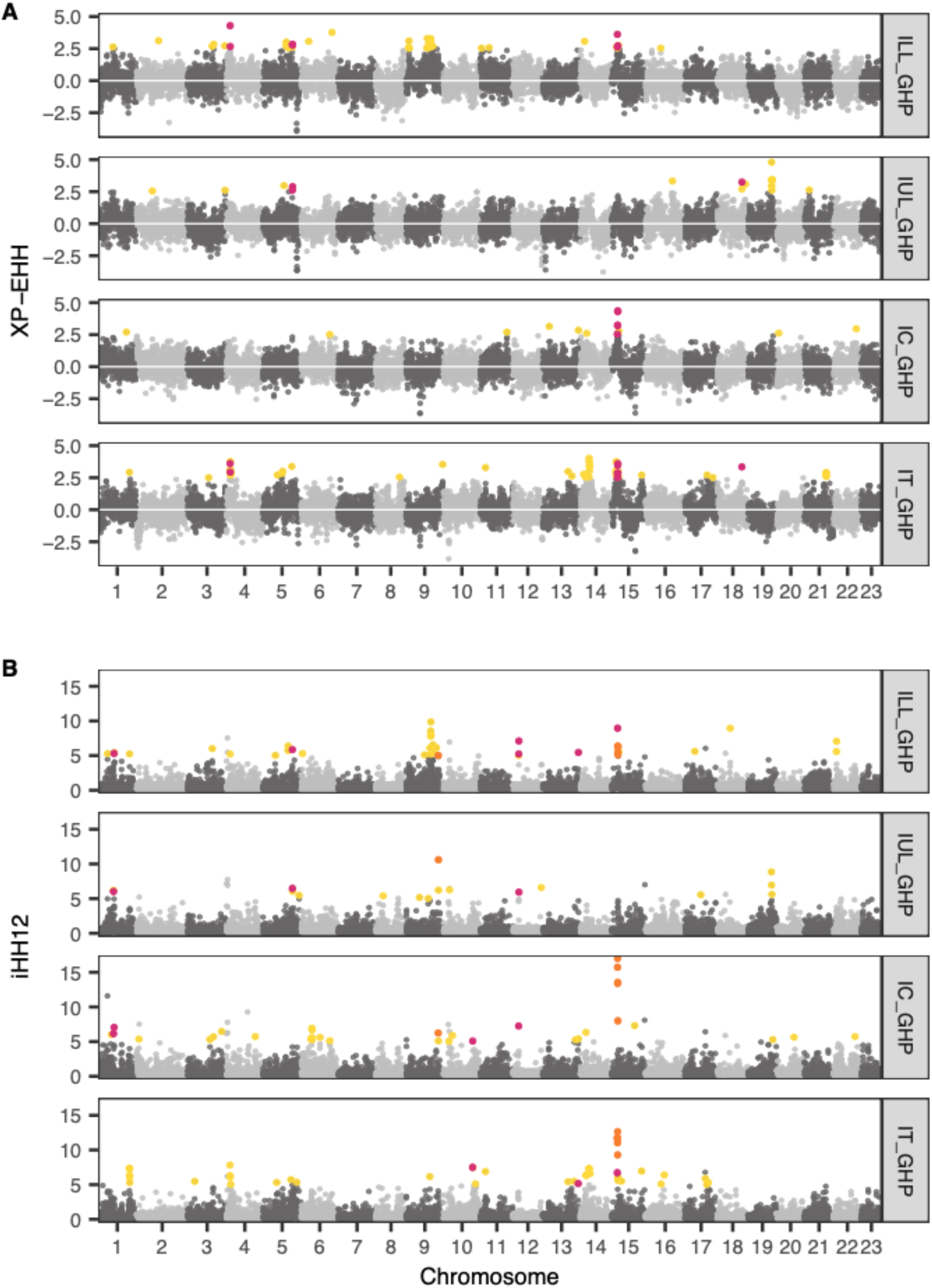
Genome-wide values of A) XP-EHH and B) iHH12 for each introduced population. Yellow points indicate unique outlier windows among the populations, pink points show overlapping outlier windows in two of the four populations, and orange points indicate windows overlapping in three of the four comparisons, there was no overlap among all four comparisons. Outliers are defined as windows with XP-EHH values > 2.5 and iHH12 values > 5.

While we found no overlap in outliers for either statistic among all 4 populations, a ~5Mb region on chromosome 15 overlapped in both statistics in 3 populations and had the highest iHH12 scores in two of the populations (figure 2). For XP-EHH, only pairwise overlaps were observed but overlapping outliers for IC, IT, and ILL pairs all involved a small region on chromosome 15 (5100000 - 5625000 bp) (table S14). Among the pairwise overlaps, IC-IT and ILL-IT overlapped more often (N=4, for both), followed by ILL-IUL (N=2), with no overlap between IUL and IC (table S14). Similarly, for iHH12,there was little overlap in outliers, and no outlier windows were common to all four comparisons. However, among the three-way overlapping outliers, ILL-IC-IT overlapped the most (N=5, expected N = <1, table S15) and all overlapping windows were located consecutively on chromosome 15 (5,175,001-5,550,000 bp). ILL-IUL-IC had two overlapping windows, one on chromosome 9 (28,500,001-28,575,000 bp) and one on an unplaced scaffold 000083F_0.2 (150,001-225,000 bp). The remaining three-way comparisons had no overlapping outlier windows. Among the pairwise overlapping outliers, comparisons including IUL showed fewer overlapping outlier windows than the other comparisons. ILL-IC and ILL-IT had the most overlapping windows (N=9 for both), followed by IC-IT (N=6), IUL-IC (N=5), ILL-IUL (N=4) and no overlap between IUL and IT. Taken together, shared signals of selection are not driven by year or canopy thinning. Finally, a region on chromosome 15 is a strong candidate for selection in three of the four introduced populations.

### Allele frequencies changed in parallel fashion in the introduced populations in a few candidate loci

Our multivariate approach revealed that the four populations exhibited parallel changes in allele frequency in both previously identified candidate loci and new loci. We identified 31 outlier windows spread over 12 of the 23 chromosomes on the first eigenvector, using the 99.9% confidence interval cut-off from the randomised allele frequency matrices (figure 3a, table S16). Eight of these windows occurred in the candidate region of convergent evolution on chromosome 15 (5,066,474 bp to 5,456,741 bp). Another cluster of eight outlier windows was on chromosome 8, covering the region between 24,649,741 bp and 25,029,959 bp (figure S3), which was not identified previously. The remaining outlier windows are distributed across the genome. In all 31 outlier windows, all populations had eigenvector loadings of the same sign and similar size (table S16), suggesting all four populations experienced parallel allele frequency changes along the same axis and in the same direction.

**Figure 3.**
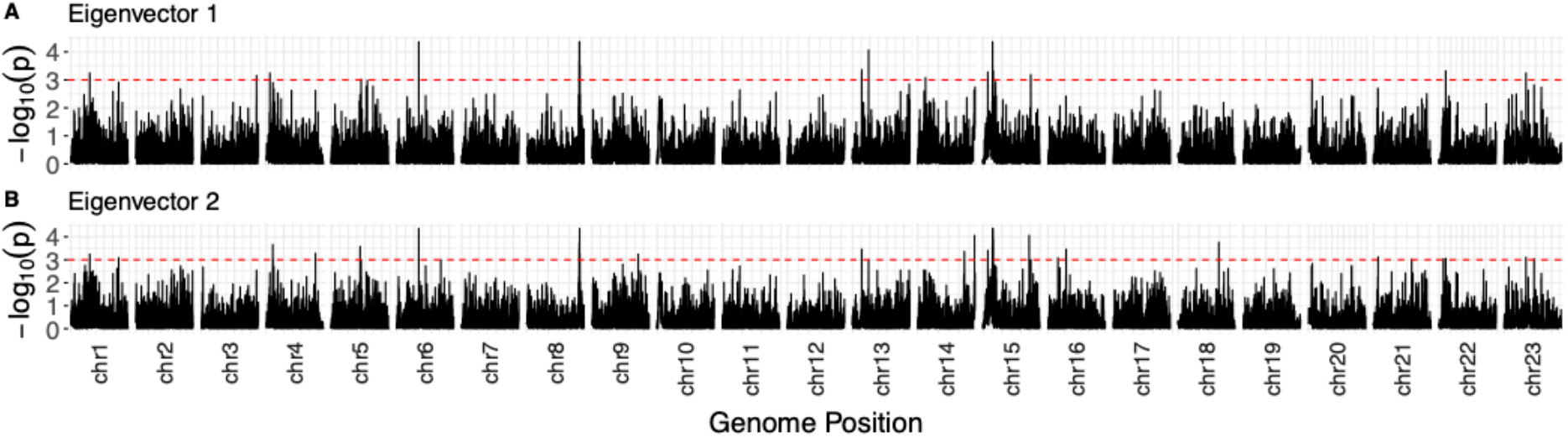
Outlier windows on (A) eigenvector 1 and (B) eigenvector 2. Red lines indicate the 99.9% confidence interval.

On eigenvector 2, we identified 30 additional outlier windows, also spread across 12 chromosomes (figure 3b, table S17). Seven of these windows occurred in our chromosome 15 candidate region (4,981,720 - 5,396,171 bp). For the outlier windows on eigenvector 2, the eigenvalues on eigenvector 1 were not significant on their own, but the sum of the eigenvalues of eigenvector 1 and 2 together was greater than expected for the sum of null eigenvalues 1 and 2. This suggests there is some residual variance that cannot be explained by the first axis alone. This provides evidence that the whole region is experiencing parallel shifts away from the GHP source genotype towards a new optimum LP genotype (eigenvector 1), but that within this newly adapted genotype there are multiple fit optima (the residual variance on eigenvector 2). This could be attributable to the presence of multiple sub-haplotypes within the parallel LP haplotype of eigenvector 1. In the convergent region on chromosome 15, we found that IUL consistently captured some residual variance; this suggests it is experiencing different allele frequency changes at a subset of sites compared to the other three populations (table S17).

### There were convergent signals of selection in all four populations in a candidate region on chromosome 15

Closer examination of allele frequency change between the GHP source and introduction populations confirmed parallel evolution at our candidate loci. The allele frequency changes of SNPs in outlier windows on chromosome 15 were broadly located along the diagonal when comparing allele frequency change for all pairwise comparisons (especially for the iHH12 regions), and they were more strongly correlated to each other than an equal sized random sample of SNPs were (p<0.0001 for all comparisons, co-correlation analysis (Diedenhofen and Musch 2015)). This is also true for pairwise comparisons with IUL, for which we found no outlier regions on chromosome 15 (figure 4a&b), confirming that all four populations are evolving along the same axis in this region.

**Figure 4.**
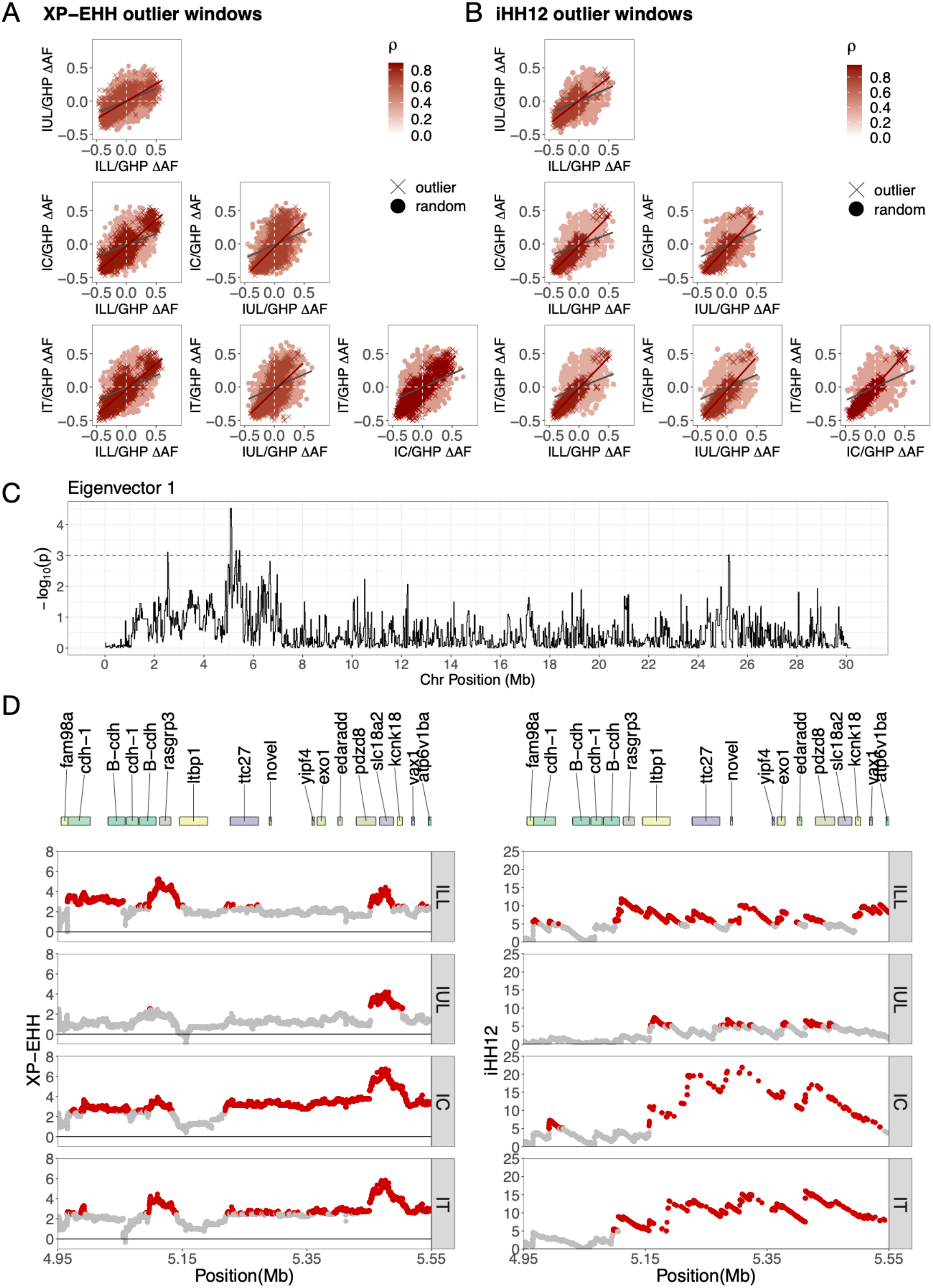
Allele frequency changes in the outlier windows on chromosome 15 for XP-EHH (A) and iHH12 (B). Crosses are allele frequency changes of SNPs in the region’s outlier windows, closed circles are allele frequency changes of a randomly drawn set of SNPs of equal size. Eigenvalues for eigenvector 1 on chromosome 15 (C), yellow points indicate windows above the 99.9% quantile. (D) Per SNP values of XP-EHH and iHH12 for the region on chromosome 15 from 4,950,000 to 5,555,000 bp (red box in C), red points indicate SNPs above the outlier cut-off values for XP-EHH and iHH12, respectively. Position and names of genes are located at the top of the plot.

This region harbours possible candidate genes involved in adaptation to the LP environment. Overlapping the outliers of eigenvector 1 and XP-EHH on chromosome 15 highlighted a region from 5,066,474 to 5,456,741 bp (figure 4c&d), and intersecting the eigenvector 1 outliers with iHH12 outlier windows revealed a similar overlapping region from 5,091,482 bp to 5,456,741 bp (figure 4d). The entire region contains eleven candidate genes (table S18). Among these are two genes encoding cadherin proteins (cadherin-1 and B-cadherin). Genes in this family have been implicated in regulating pigment cell migration, and are involved in cell-cell adhesion interactions (Fukuzawa and Obika 1995; Nishimura *et al.* 1999). Another candidate gene in this region is solute carrier 18 member A2 (*slc18a2*, figure 4c&d), which is involved in various growth and body-size phenotypes in mice, including head morphology (Bult *et al.* 2019; Stelzer *et al.* 2016). Guppies from HP and LP environments have previously been shown to have different head morphologies, where LP guppies had shorter and rounder heads than HP guppies (Torres-Dowdall *et al.* 2012). Taking all the evidence together, this region on chromosome 15 shows strong evidence for selection in at least three of the four populations and experienced parallel allele frequency changes among all four populations.

## DISCUSSION

Our four rapidly evolving, experimental populations of guppies reveal only small genome-wide differentiation from to their HP ancestors. Despite these small genome-wide changes, we uncovered strong local signals of selection in all four populations at individual loci. Using a combination of haplotype genome scans and a newly developed multivariate approach, we found evidence from the ancestor and parallel change among the four descendants at candidate loci. Our multivariate approach revealed more subtle parallel changes in allele frequency compared to the genome scans. Combining the evidence, we found a region under strong selection on chromosome 15 in three of the four populations and the multivariate approach showed all four populations are evolving in parallel in this region.

We also found little evidence for bottlenecks and inbreeding in three of the four populations (ILL, IUL and IC), whereas one population (IT) exhibited patterns consistent with inbreeding and bottlenecks in the recent past. This is in agreement with past census data from these populations and the population crash seen in IT during the first year of the introduction (Reznick *et al.* 2019).

A region on chromosome 15 (positioned at 5Mb), in which we identified several overlapping outliers and parallel change among all populations, is a strong candidate for convergent evolution. This region has previously been identified as a candidate for convergent evolution in natural populations and long-term experimental populations, in a RADSeq study (specifically in the Oropuche, Arima, and introduction Aripo HP-LP pairs) (Fraser *et al.* 2015). Further, using whole-genome sequencing, this same region was found to be a candidate of selection in the Tacarigua and Oropuche population pairs and a region slightly upstream in the Aripo and Tacarigua pairs (Whiting *et al.* 2020). Finally, the cadherin signaling pathway was enriched for signatures of selection in all paired comparisons of HP and LP populations studied (Whiting *et al.* 2020). Cadherin family genes are known to be involved in pigment cell migration and other cell-cell adhesion processes (Fukuzawa and Obika 1995; Nishimura *et al.* 1999), and could thus be involved in the colour differences between HP and LP males. Further studies, such as gene knockout experiments, could help identify the exact role of the cadherin genes in this process.

Rapid convergent genomic adaptation, specifically at quantitative traits, is often predicted to occur through small shifts in allele frequency (Barrett and Schluter 2008), which are difficult to detect using most available methods. The novel multivariate allele frequency change analysis described here can identify parallel shifts in allele frequency rather than absolute changes in allele frequency at individual loci. Using this method, we found 31 windows experiencing parallel changes in allele frequency among the four populations.

Most of these 31 windows lacked a strong haplotype signal in the genome scan measures, which can be explained in several ways. First, it is possible the variance on this axis was constrained for reasons other than positive selection. For example, background selection (BGS) could cause correlated differentiation landscapes among the populations, which results in a constrained axis of allele frequency changes (Burri 2017). However, it is unlikely that BGS would produce the fully parallel signals we observed here. Using simulations, others (Matthey-Doret and Whitlock 2019; Stankowski *et al.* 2019) have shown that when divergence times are short, BGS is unlikely to generate correlated differentiation landscapes. Other processes, such as positive selection, are the more likely cause of these patterns. Second, the strength of selection may have been relatively weak, causing time of fixation for a beneficial allele to be considerably longer. These weakly selected alleles would not have had time to generate extreme frequency differences in the introduced populations (Coop *et al.* 2009). However, because we detected parallel allele frequency change in all four populations, its actual variance relative to the rest of the genome is less important. Finally, many of the phenotypic traits under selection in LP are likely to have polygenic control. If this is the case, then simultaneous selection on standing genetic variation at many loci would cause subtle shifts of allele frequencies at individual loci (Pritchard *et al.* 2010). These factors would result in small parallel shifts in allele frequencies rather than complete sweeps at single loci.

It is widely appreciated that demographic changes in population size can create patterns that resemble the consequences of selection. To explore the genomic signatures of these events, we used genome-wide patterns of variation and haplotype statistics. Compared to the other introduced populations, IT had the lowest levels of expected heterozygosity and nucleotide diversity, the greatest increase in Tajima’s D, and more and longer ROH, all indicators of a recent bottleneck (Smith and Haigh 2007; Tajima 1989). Reznick et al. (2019) observed a severe crash in the population density of IT during the first year, and two other studies (Dowdall *et al.* 2012; Fitzpatrick *et al.* 2014) found IT had a higher mortality rate than IC in the first year, which they attributed to stronger influence of seasonal changes and indicators of disease in IT. Together, these results suggest that the bottleneck we detected in IT probably occurred in the first year after the introduction, as opposed to a founding bottleneck followed by population growth. It is satisfying that we were able to detect this bottleneck in the genome-wide data by combining site frequency tests with haplotype homozygosity tests. Our ability to detect the consequences of a known bottleneck highlights the validity of these methods in recently established populations, and that it can be a useful approach even for studies without available census data.

It could be expected that as a result of the bottleneck, IT might have experienced a different selection pressure compared to the other populations (e.g. less effective because of chance loss of alleles), or selection on a different set of variants (e.g. strong immune responses to overcome the increased incidence of disease reported). However, we find no evidence for this. Average allele frequency changes were similar across the four populations. Further, in our overlapping outlier analysis, comparisons including IT consistently had high numbers of overlapping outlier windows, suggesting IT was not experiencing a different selection pressure than the other introductions.

Rather, comparisons with IUL had the lowest overlapping windows in the outlier analysis and increased levels of genome-wide heterozygosity. This population was one of the sites initiated with low density (IUL and ILL) and also had a thinned canopy. As a result, IUL had the largest population sizes by far during the dry season, but also had the strongest population declines (Reznick *et al.* 2019). Combined, this could result in chance allele frequency changes associated with the period of population growth and could reduce our ability to detect signatures of selection, i.e. reduced diversity.

The onset of selection coincided with the populations’ attaining peak densities (Reznick *et al.* 2019), the timing of which differed between the pairs of populations because of their different initial densities. Populations with low initial densities (ILL and IUL) took three years to attain peak density, whereas the high initial densities (IC and IT) took only two years to attain peak density and hence started to evolve after just two years. We find evidence of these differences in population growth and contraction at the genomic scale. The lower density populations, ILL and IUL, saw a slight increase in expected heterozygosity and nucleotide diversity and a reduced Tajima’s D, which is indicative of a population that experienced rapid population expansion (Ortego *et al.* 2007; Zenger *et al.* 2003). In the higher density populations, IC and IT, we instead found reduced expected heterozygosity and nucleotide diversity, and increased Tajima’s D. Both are signals of reduced population growth.

Additionally, we show that differences in initial population density also led to IC and IT having more SNPs with unchanged allele frequencies, likely because there was less time for SNPs to randomly drift away from GHP allele frequencies. We further find that IC and IT have more fixed SNPs that were the minor allele in GHP, as well as having more overlapping outlier windows in the genome scans, suggesting selection had a stronger effect in the high-density populations compared to the low-density populations.

In conclusion, we found strong signals of selection in all four populations, and a region on chromosome 15 contained overlapping windows with strong signals of selection in three of the four populations. Our novel multivariate analysis of allele frequencies revealed evidence of subtle parallel changes in allele frequency in this region. This suggests this method has the potential to detect convergent evolution in rapidly evolving populations, as well as identifying signatures of polygenic selection with small changes at many loci simultaneously.

## ACKNOWLEDGEMENTS

We thank Tim Coulson and the University of Exeter’s Population Genetics group for helpful discussion on the manuscript. We also acknowledge high performing computing (HPC) ISCA server at the University of Exeter. This work was supported by the Max Planck Society (DW), EU Research Council grant (GuppyCon 758382) (BAF, JRW), NERC grant (NE/P013074/1) (JRP), University of Sussex and University of Exeter (MJvdZ) and the National Science Foundation USA (DEB-0623632EF, DEB-0808039, DEB-1258231, DEB-1556884) to DNR, JT, RB. The authors declare no conflicts of interest.

## AUTHOR CONTRIBUTION

BAF, DNR, RB, JT, and DW conceived of the study. DNR, RB, and JT conducted the initial introduction experiment. Genomic work and analysis was completed by MJvdZ, JRW, JRP, and BAF. Writing was done by all authors.

## DATA ACCESSIBILITY

All scripts are available on Github repository: https://github.com/josieparis/gatk-snp-calling https://github.com/mvdzee/Rapid_genomic_adaptation_guppies ENA accession numbers for sequences: PRJEB42705 (all introduction populations) and PRJEB10680 (GHP).

## SUPPLEMENTARY METHODS

### Genome sequencing and SNP calling

Fish were stored in 95% ethanol at −20C prior to DNA extraction. Genomic DNA was extracted from caudal peduncle tissue using the Qiagen DNeasy Blood and tissue kit (QIAGEN; Hilden, Germany). Whole genome sequencing libraries were prepared following the Illumina TruSeq DNA sample preparation guide with approximately 250bp insert size. Eighty-six samples were sequenced by multiplexing 6-10 individuals per lane on an Illumina HiSeq 2000 and 3000.The remaining eight samples were sequenced using the Illumina HiSeq 4000 with a 150bp paired-end read metric.

Quality of paired-end reads was assessed with FastQC (Andrews 2010) and adapters and low-quality bases removed with TrimGalore! (Krueger 2012). Reads were aligned to the long-read, updated guppy reference genome (Fraser *et al.* 2020) using BWA-mem (v0.7.17)(Li and Durbin 2009). Read groups were added and duplicate reads were removed before merging to produce final bams using picard v 2.06. Read quality was recalibrated using variants generated from high-coverage, PCR-free sequencing data (Fraser *et al.* 2020). GVCFs were obtained using GATK’s (v4.0.5.1) HaplotypeCaller and combined with GenomicsDBImport before genotyping with GenotypeGVCFs. Resulting variants were filtered on the basis of QD<2.0, FS>60.0, MQ<40.0, HaplotypeScore>13.0 and MappingQualityRankSum < −12.5 according to GATK best practices, and only bi-allelic sites were retained. SNPs were removed if they had a depth <5x and if missing in 50% of individuals. Finally, the population files were merged again and filtered for a minor allele frequency of >0.01. The final VCF file contained 6,510,265 SNPs.

For the haplotype analyses, we followed the double-phasing protocol as described in (Malinsky *et al.* 2018): population VCF files were first phased per chromosome using Beagle (v5.0) (Browning and Browning 2007), followed by a second round of phasing with Shapeit2 (v2.r904) (Delaneau *et al.* 2011). Shapeit2 has an increased accuracy compared to beagle (Delaneau *et al.* 2011), but does not accept missing data. Therefore, we use beagle, which does accept missing data but has a high switch rate error (Delaneau *et al.* 2011), to create pre-phased VCF files that can be used as input for shapeit2.

### Runs of homozygosity

Runs of homozygosity (ROH) were calculated for each individual with a sliding window approach with 50 SNPs per window using Plink. To minimize the detection of ROH that could occur by chance, the minimum number of SNPs needed to constitute a ROH (l) was estimated using the method proposed by (Lencz *et al.* 2007). Additionally, each run had to be at least 500 kb long to exclude short, common ROH present in all individuals and populations. Finally, at most 1 heterozygous site per window was allowed. Runs of homozygosity were estimated for each individual separately, and resulting ROH were binned into four categories: 0.5-0.75 Mb, 0.75-1.0 Mb, 1.0-1.5 Mb, and >1.5 Mb. To calculate the genomic inbreeding coefficient F_ROH_, we used:

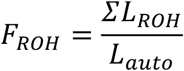

Where L_ROH_ is the total length of all of an individual’s ROH above a specified length threshold and L_auto_ is the length of the autosomal genome.

### Genome scans

Both XP-EHH and iHH12 were calculated using Selscan (v1.2.0a) (Szpiech and Hernandez 2014), using the default settings, except for a MAF filter of 0.01. The results were then normalised over all chromosomes using the script provided by Selscan. XP-EHH outliers were identified by a value of XP-EHH > 2.5, and iHH12 outliers were identified as those windows with an absolute value of iHH12 > 5 in the introduced populations and an absolute value of <5 in the GHP source.

The number of overlapping outliers among populations per measure was calculated using the R package SuperExactTest (Wang *et al.* 2015). This package also calculates the expected number of overlapping windows for each set based on a hypergeometric distribution.

To assess the extent of parallel allele frequency changes in the outlier region, we extracted allele frequency changes per SNP (ΔAF) among the SNPs in the XP-EHH and iHH12 outlier windows and made pairwise plots of ΔAF. We compared these values to values of ΔAF to a pool of equal size drawn randomly across the genome. Finally, we obtained gene annotations for the outlier region by aligning the region to the previously published female genome (Künstner *et al.* 2016) and extracting guppy genes using Ensembl’s BioMart (release 102) (Howe *et al.* 2021).

### Multivariate analysis of parallel allele frequency changes

Briefly, we divided each chromosome into non-overlapping windows of 200 SNPs, and calculated a matrix of normalised allele frequency changes per window:

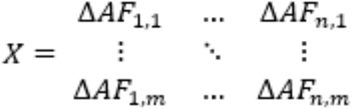

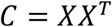

Eigen decomposition of each C matrix produces a distribution of eigenvalues; distributions with excessive variance explained by the first eigenvector suggest a shared direction of allele frequency changes among all population pairs through multivariate space. This direction of change could be parallel (along the same axis in the same direction) or anti-parallel (along the same axis in different directions). We created a null-distribution to determine which windows show extreme eigenvalues. We randomly sampled windows of an equal number of SNPs (N=200) along the genome, and allowed for a random start position for windows. For each null permutation, individual IDs were shuffled and allele frequency vectors re-calculated between random pairs. We applied the same transformations described above to each window and ran 1000 permutations. Windows were identified as outliers if they had an eigenvalue above the genome-wide 99.9% quantile of the null-distribution.

Finally, we investigated where the allele frequency changes are parallel and where they are anti-parallel by examining the loading of each population on the first eigenvector. Lineages with the same sign loading can be interpreted as evolving in the same direction, and lineages with opposite signs are evolving anti-parallel.

## SUPPLEMENTARY FIGURES

**Figure S1:**
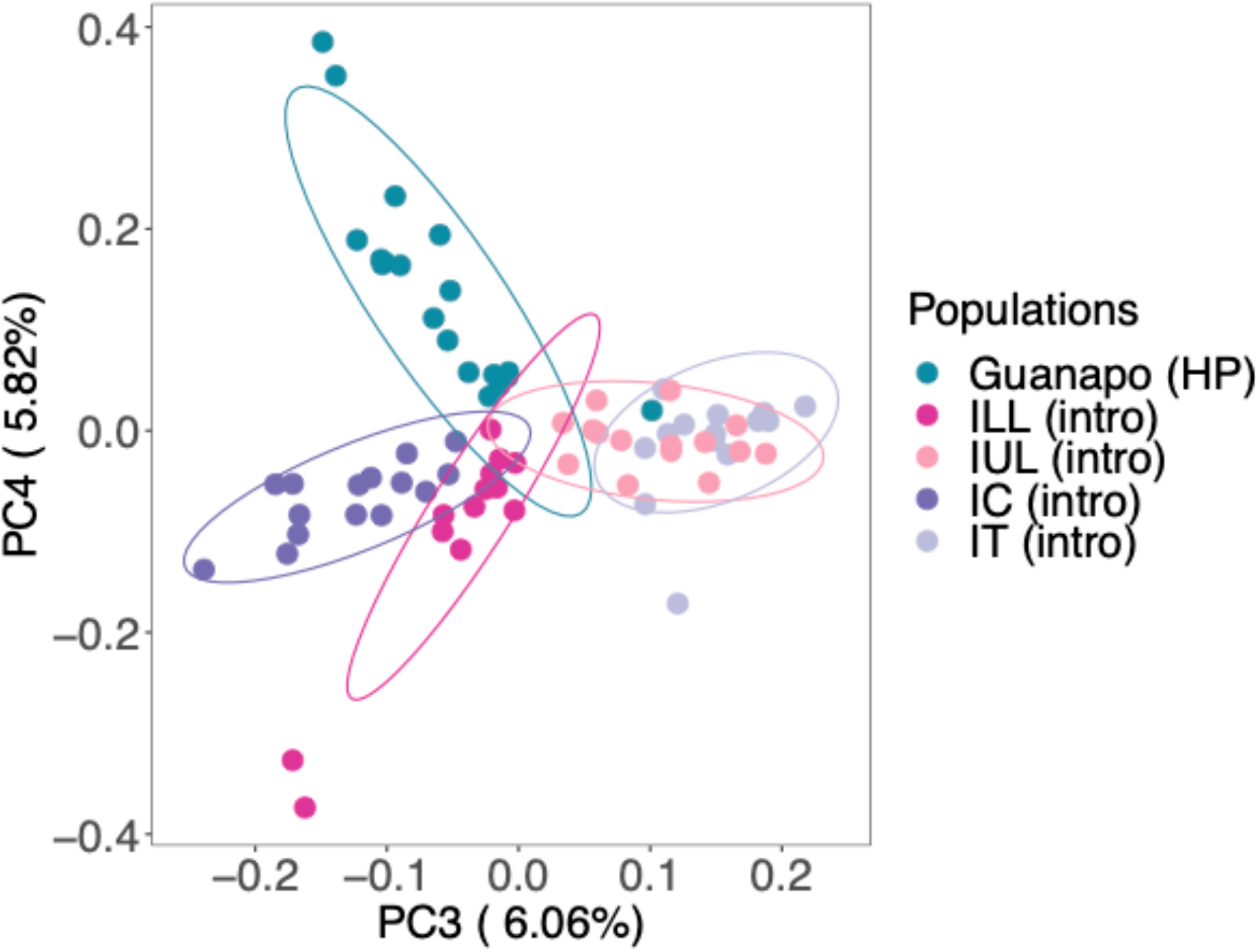
PC3 and PC4 with populations coloured according to river.

**Figure S2.**
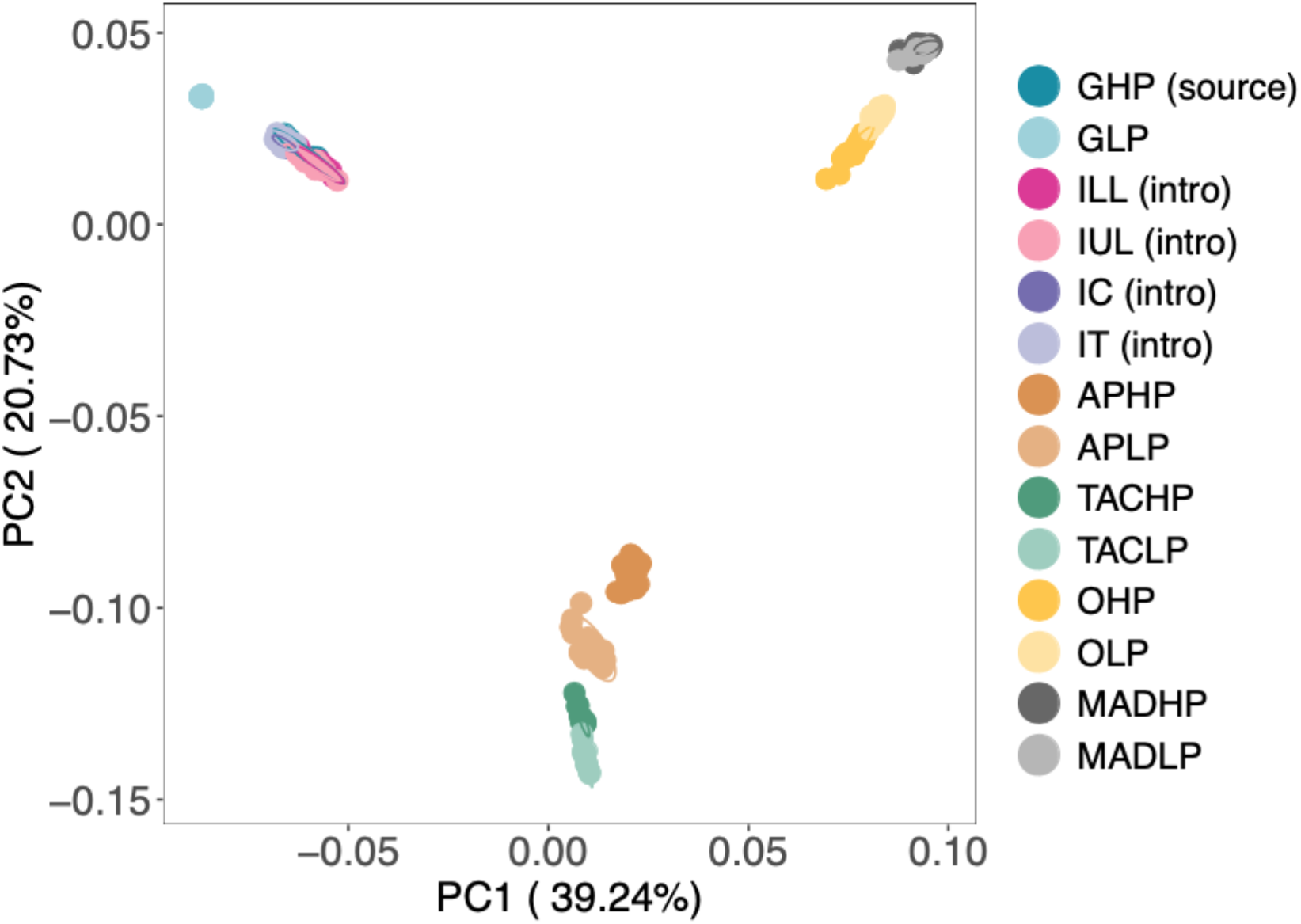
Principal component analysis with natural populations described in (Whiting et al., 2020) illustrating the lack of population structure between GHP and the introduced populations.

**Figure S3.**
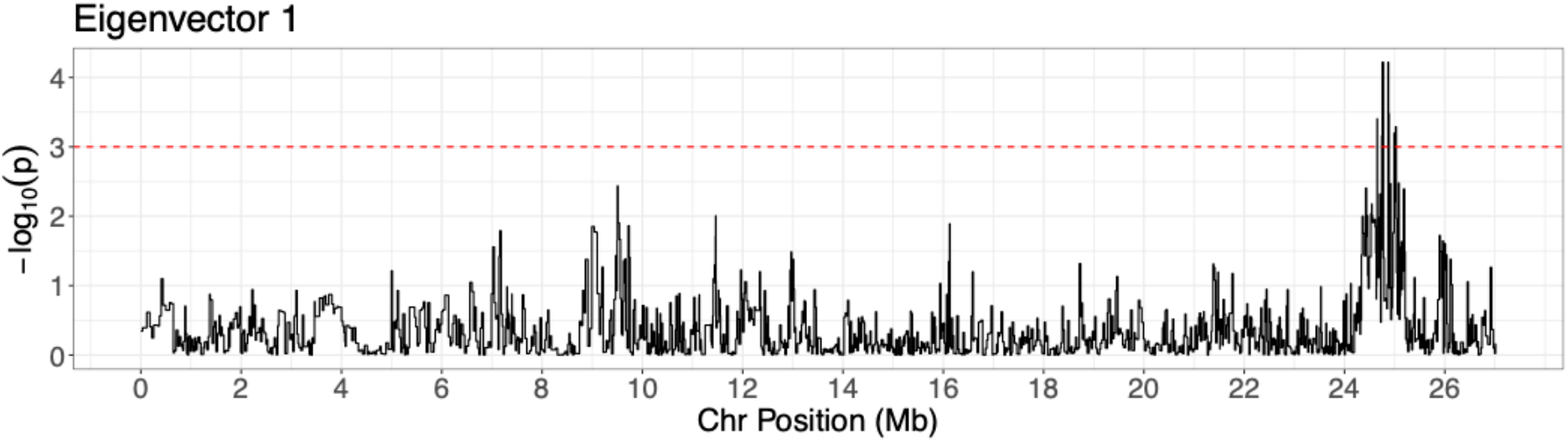
Eigenvalues for eigenvector 1 on chromosome 8, yellow points indicate windows above the 99.9% quantile. Red box indicates the region with consecutive outlier windows

## REFERENCES

Arendt, J.D., Reznick, D.N. & López-Sepulcre, A. (2014). Replicated origin of female-biased adult sex ratio in introduced populations of the trinidadian guppy (*Poecilia reticulata*). Evolution 68:2343–2356.

Barrett, R.D.H. & Schluter, D. (2008). Adaptation from standing genetic variation. Trends Ecol. Evol. 23:38–44.

Bult, C.J., Blake, J.A., Smith, C.L., Kadin, J.A., Richardson, J.E. & Mouse Genome Database Group (2019). Mouse Genome Database (MGD) 2019. Nucleic Acids Res. 47:D801–D806.

Burri, R. (2017). Interpreting differentiation landscapes in the light of long-term linked selection. Evolution Letters 1:118–131.

Chang, C.C., Chow, C.C., Tellier, L.C., Vattikuti, S., Purcell, S.M. & Lee, J.J. (2015). Second-generation PLINK: rising to the challenge of larger and richer datasets. Gigascience 4:7.

Coop, G., Pickrell, J.K., Novembre, J., Kudaravalli, S., Li, J., Absher, D., Myers, R.M., Cavalli-Sforza, L.L., Feldman, M.W. & Pritchard, J.K. (2009). The role of geography in human adaptation. PLoS Genet. 5:e1000500.

Danecek, P., Auton, A., Abecasis, G., Albers, C.A., Banks, E., DePristo, M.A., Handsaker, R.E., Lunter, G., Marth, G.T., Sherry, S.T., et al. (2011). The variant call format and VCFtools. Bioinformatics 27:2156–2158.

De Lisle, S.P. & Bolnick, D.I. (2020). A multivariate view of parallel evolution. Evolution 74:1466–1481.

Diedenhofen, B. & Musch, J. (2015). cocor: a comprehensive solution for the statistical comparison of correlations. PLoS One 10:e0121945.

Dowdall, J.T., Handelsman, C.A., Ruell, E.W., Auer, S.K., Reznick, D.N. & Ghalambor, C.K. (2012). Fine-scale local adaptation in life histories along a continuous environmental gradient in Trinidadian guppies. Functional Ecology 26:616–627.

Endler, J.A. (1980). Natural Selection on color patterns in Poecilia reticulata. Evolution 34:76–91.

Endler, J.A. & Houde, A.E. (1995). Geographic variation in female preferences for male traits in *Poecilia reticulata*. Evolution 49:456–468.

Fitzpatrick, S.W., Torres-Dowdall, J., Reznick, D.N., Ghalambor, C.K. & Funk, W.C. (2014). Parallelism isn’t perfect: could disease and flooding drive a life-history anomaly in Trinidadian guppies? Am. Nat. 183:290–300.

Fraser, B.A., Künstner, A., Reznick, D.N., Dreyer, C. & Weigel, D. (2015). Population genomics of natural and experimental populations of guppies (Poecilia reticulata). Mol. Ecol. 24:389–408.

Fraser, B.A., Whiting, J.R., Paris, J.R., Weadick, C.J., Parsons, P.J., Charlesworth, D., Bergero, R., Bemm, F., Hoffmann, M., Kottler, V.A., et al. (2020). Improved Reference Genome Uncovers Novel Sex-Linked Regions in the Guppy (Poecilia reticulata). Genome Biol. Evol. 12:1789–1805.

Fukuzawa, T. & Obika, M. (1995). N-CAM and N-cadherin are specifically expressed in xanthophores, but not in the other types of pigment cells, melanophores, and iridiphores. Pigment Cell Res. 8:1–9.

Garud, N.R., Messer, P.W., Buzbas, E.O. & Petrov, D.A. (2015). Recent selective sweeps in North American Drosophila melanogaster show signatures of soft sweeps. PLoS Genet. 11:e1005004.

Gillespie, J.H. (1994). The Causes of Molecular Evolution (Oxford University Press on Demand).

Grant, P.R. & Grant, R. B. (2020). How and Why Species Multiply: The Radiation of Darwin’s Finches (Princeton University Press).

Handelsman, C.A., Broder, E.D., Dalton, C.M., Ruell, E.W., Myrick, C.A., Reznick, D.N. & Ghalambor, C.K. (2013). Predator-induced phenotypic plasticity in metabolism and rate of growth: rapid adaptation to a novel environment. Integr. Comp. Biol. 53:975–988.

Hermisson, J. & Pennings, P.S. (2005). Soft sweeps: molecular population genetics of adaptation from standing genetic variation. Genetics 169:2335–2352.

Kemp, D.J., Reznick, D.N., Grether, G.F. & Endler, J.A. (2009). Predicting the direction of ornament evolution in Trinidadian guppies (Poecilia reticulata). Proc. Biol. Sci. 276:4335–4343.

Kemp, D.J., Batistic, F.-K. & Reznick, D.N. (2018). Predictable adaptive trajectories of sexual coloration in the wild: Evidence from replicate experimental guppy populations. Evolution 72:2462–2477.

Kohler, T.J., Heatherly, T.N., El-Sabaawi, R.W., Zandonà, E., Marshall, M.C., Flecker, A.S., Pringle, C.M., Reznick, D.N. & Thomas, S.A. (2012). Flow, nutrients, and light availability influence Neotropical epilithon biomass and stoichiometry. Freshwater Science 31:1019–1034.

Lee, K.M. & Coop, G. (2017). Distinguishing Among Modes of Convergent Adaptation Using Population Genomic Data. Genetics 207:1591–1619.

Losos, J.B. (2011). Lizards in an Evolutionary Tree: Ecology and Adaptive Radiation of Anoles (Univ of California Press).

Lowry, D.B., Hoban, S., Kelley, J.L., Lotterhos, K.E., Reed, L.K., Antolin, M.F. & Storfer, A. (2017). Breaking RAD: an evaluation of the utility of restriction site-associated DNA sequencing for genome scans of adaptation. Molecular Ecology Resources 17:142–152.

Magurran, A.E. (2005). Evolutionary Ecology: The Trinidadian Guppy (Oxford University Press on Demand).

Malinsky, M., Svardal, H., Tyers, A.M., Miska, E.A., Genner, M.J., Turner, G.F. & Durbin, R. (2018). Whole-genome sequences of Malawi cichlids reveal multiple radiations interconnected by gene flow. Nat Ecol Evol 2:1940–1955.

Matthey-Doret, R. & Whitlock, M.C. (2019). Background selection and FST : Consequences for detecting local adaptation. Mol. Ecol. 28:3902–3914.

Nishimura, E.K., Yoshida, H., Kunisada, T. & Nishikawa, S.-I. (1999). Regulation of E- and P-Cadherin Expression Correlated with Melanocyte Migration and Diversification. Developmental Biology 215:155–166.

Ortego, J., Aparicio, J.M., Calabuig, G. & Cordero, P.J. (2007). Increase of heterozygosity in a growing population of lesser kestrels. Biol. Lett. 3:585–588.

Pascoal, S., Cezard, T., Eik-Nes, A., Gharbi, K., Majewska, J., Payne, E., Ritchie, M.G., Zuk, M. & Bailey, N.W. (2014). Rapid convergent evolution in wild crickets. Curr. Biol. 24:1369–1374.

Pfeifer, B., Wittelsbürger, U., Ramos-Onsins, S.E. & Lercher, M.J. (2014). PopGenome: an efficient Swiss army knife for population genomic analyses in R. Mol. Biol. Evol. 31:1929–1936.

Pritchard, J.K., Pickrell, J.K. & Coop, G. (2010). The genetics of human adaptation: hard sweeps, soft sweeps, and polygenic adaptation. Curr. Biol. 20:R208–R215.

Reznick, D.N. & Endler, J.A. (1982). The impact of predation on life history evolution in Trinidadian guppies (Poecilia reticulata). Evolution 36:160–177.

Reznick, D.N. & Bryga, H.A. (1996). Life-History Evolution in Guppies (Poecilia reticulata: Poeciliidae). V. Genetic Basis of Parallelism in Life Histories. The Am. Nat. 147:339–359.

Reznick, D.N., Bassar, R.D., Handelsman, C.A., Ghalambor, C.K., Arendt, J., Coulson, T., Potter, T., Ruell, E.W., Torres-Dowdall, J., Bentzen, P., et al. (2019). Eco-Evolutionary Feedbacks Predict the Time Course of Rapid Life-History Evolution. Am. Nat. 194:671–692.

Sabeti, P.C., Varilly, P., Fry, B., Lohmueller, J., Hostetter, E., Cotsapas, C., Xie, X., Byrne, E.H., McCarroll, S.A., Gaudet, R., et al. (2007). Genome-wide detection and characterization of positive selection in human populations. Nature 449:913–918.

Smith, J.M. & Haigh, J. (2007). The hitch-hiking effect of a favourable gene. Genet. Res. 89:391–403.

Stankowski, S., Chase, M.A., Fuiten, A.M., Rodrigues, M.F., Ralph, P.L. & Streisfeld, M.A. (2019). Widespread selection and gene flow shape the genomic landscape during a radiation of monkeyflowers. PLoS Biol. 17:e3000391.

Stelzer, G., Rosen, N., Plaschkes, I., Zimmerman, S., Twik, M., Fishilevich, S., Stein, T.I., Nudel, R., Lieder, I., Mazor, Y., et al. (2016). The GeneCards Suite: From Gene Data Mining to Disease Genome Sequence Analyses. Curr. Protoc. Bioinformatics 54:1.30.1–1.30.33.

Tajima, F. (1989). The effect of change in population size on DNA polymorphism. Genetics 123:597–601.

Tiffin, P. & Ross-Ibarra, J. (2014). Advances and limits of using population genetics to understand local adaptation. Trends Ecol. Evol. 29:673–680.

Torres, R., Szpiech, Z.A. & Hernandez, R.D. (2018). Human demographic history has amplified the effects of background selection across the genome. PLoS Genet. 14:e1007387.

Torres-Dowdall, J., Handelsman, C.A., Reznick, D.N. & Ghalambor, C.K. (2012). Local adaptation and the evolution of phenotypic plasticity in Trinidadian guppies (Poecilia reticulata). Evolution 66:3432–3443.

Travis, J., Reznick, D., Bassar, R.D., López-Sepulcre, A., Ferriere, R. & Coulson, T. (2014). Do Eco-Evo Feedbacks Help Us Understand Nature? Answers From Studies of the Trinidadian Guppy. Eco-Evolutionary Dynamics 1–40.

Whiting, J.R., Paris, J.R., van der Zee, M.J., Parsons, P.J., Weigel, D. & Fraser, B.A. (2020). Drainage-structuring of ancestral variation and a common functional pathway shape limited genomic convergence in natural high- and low-predation guppies.

Zenger, K.R., Richardson, B.J. & Vachot-Griffin, A.-M. (2003). A rapid population expansion retains genetic diversity within European rabbits in Australia. Mol. Ecol. 12:789–794.

## REFERENCES

Andrews, S. (2010). FastQC: a quality control tool for high throughput sequence data.

Browning, S.R. & Browning, B.L. (2007). Rapid and accurate haplotype phasing and missing-data inference for whole-genome association studies by use of localized haplotype clustering. Am. J. Hum. Genet. 81:1084–1097.

Delaneau, O., Marchini, J. & Zagury, J.-F. (2011). A linear complexity phasing method for thousands of genomes. Nat. Methods 9:179–181.

Howe, K.L., Achuthan, P., Allen, J., Allen, J., Alvarez-Jarreta, J., Amode, M.R., Armean, I.M., Azov, A.G., Bennett, R., Bhai, J., et al. (2021). Ensembl 2021. Nucleic Acids Res. 49:D884–D891.

Krueger, F. (2012). A wrapper tool around Cutadapt and FastQC to consistently apply quality and adapter trimming to FastQ files.

Künstner, A., Hoffmann, M., Fraser, B.A., Kottler, V.A., Sharma, E., Weigel, D. & Dreyer, C. (2016). The Genome of the Trinidadian Guppy, Poecilia reticulata, and Variation in the Guanapo Population. PLoS One 11:e0169087.

Lencz, T., Lambert, C., DeRosse, P., Burdick, K.E., Morgan, T.V., Kane, J.M., Kucherlapati, R., & Malhotra, A.K. (2007). Runs of homozygosity reveal highly penetrant recessive loci in schizophrenia. Proc. Natl. Acad. Sci. U. S. A. 104:19942–19947.

Li, H., and Durbin, R. (2009). Fast and accurate short read alignment with Burrows-Wheeler transform. Bioinformatics 25:1754–1760.

Szpiech, Z.A. & Hernandez, R.D. (2014). selscan: an efficient multithreaded program to perform EHH-based scans for positive selection. Mol. Biol. Evol. 31, 2824–2827.

Wang, M., Zhao, Y., & Zhang, B. (2015). Efficient Test and Visualization of Multi-Set Intersections. Sci. Rep. 5:16923.

Whiting, J.R., Paris, J.R., van der Zee, M.J., Parsons, P.J., Weigel, D., & Fraser, B.A. (2020). Drainage-structuring of ancestral variation and a common functional pathway shape limited genomic convergence in natural high- and low-predation guppies.

